# Affiliation in mice induced by an auditory-conditioned stimulus

**DOI:** 10.64898/2026.02.11.705410

**Authors:** Wataru Ito, Alexei Morozov

## Abstract

Social affiliation is a vital survival strategy that promotes well-being across species. While innate threats consistently trigger affiliation, it remains unclear whether learned threats engage the same affiliative systems. Here, we report that an auditory conditioned stimulus induced affiliation in mice. In same-sex dyads, mice subjected to auditory fear conditioning increased proximity during presentation of the conditioned stimulus (CS). This CS-evoked affiliation was independent of freezing levels, required familiarity between partners, and depended on intact basolateral amygdala-to-ventral hippocampus inputs and oxytocin signaling. These findings suggest that neural pathways activated by learned and innate threats converge on shared social-affiliation circuitry.

## Introduction

Affiliation is a core social behavior that supports survival and well-being by enabling mating, territorial defense, thermoregulation through huddling, skill acquisition, and collective defense against predators [1-4]. Classic fear-affiliation theory, proposed by Schachter, suggests that perceived threat actively drives social contact, particularly with others facing the same danger [5,6]. Consistent with this idea, defensive affiliation is widely observed across species in response to innate threats [7-15], and laboratory studies show that predator cues can induce aggregation in rats via neuropeptide systems, including oxytocin and vasopressin [16-18]. Collectively, these observations suggest that defensive affiliation reflects an adaptive coupling between threat-processing and social-motivational systems. This coupling is particularly relevant to understanding psychiatric disorders, in which both threat-processing and social motivation are frequently disrupted. A key unanswered question is whether this coupling extends beyond innate threats to learned threat predictors. Specifically, do conditioned threat cues elicit affiliation, and if so, which circuits and molecules mediate this effect? Here, we demonstrate that an auditory conditioned stimulus triggers affiliation in fear-conditioned mice tested as dyads. This behavior requires basolateral amygdala-to-ventral hippocampus circuitry and oxytocin signaling.

## Materials and Methods

All procedures on animals were conducted in accordance with Virginia Tech IACUC-approved protocol 24-110.

### Animals

129SvEv/C57BL/6N F1 hybrids were produced in-house from breeding trios consisting of one C57BL/6N male and two 129SvEv females, weaned at postnatal day 23–28 (p23– 28), and housed as four same-sex littermates per cage [19]. Mice were tested at p75–90. Familiar dyads were assembled 7 days before testing from cagemate littermates and housed in fresh bedding that remained unchanged until testing completion. Unfamiliar dyads were assembled at the time of testing by pairing non-littermate mice that had never been co-housed.

### Fear conditioning

Each mouse was conditioned individually in standard chambers (Med Associates, St. Albans, VT) [20] using four pairings of a conditioned stimulus (CS: 30 s, 8 kHz, 80 dB tone) with an unconditioned stimulus (US: 0.5 mA, 0.5 s footshock co-terminating with the CS), presented at variable intervals (60–180 s). Testing occurred 1–2 days later in a novel context with two mice per chamber. Each test session lasted 3 minutes: 1 minute baseline (pre-CS) followed by 2 minutes of CS presentation. Videos were recorded at 4 frames per second and exported as AVI files with MJPEG compression using Freezeframe software (Actimetrics, Wilmette, IL), then converted to MP4 format using a custom Python script.

### Animal tracking and distance estimation

Using a custom Python script, we manually annotated the snout position of each animal in every video frame and recorded the pixel coordinates. Real-world X–Y coordinates of each snout’s projection onto the chamber floor were computed using a triangulation-based geometric transformation. This transformation utilized the pixel coordinates of the snout and three chamber corners, along with the known real-world coordinates of those corners on the chamber floor plane. Intermouse distances were calculated from the resulting X–Y coordinates.

### Quantification of freezing

Freezing was defined as the absence of observable movement of the body and vibrissae (except respiration) for at least 1 s [21]. Trained observers manually recorded the first and last video frames of each freezing bout using a custom Python script, generating time-series data. Freezing percentage was computed from these data using another Python script. Averaged CS-induced freezing across all groups ranged from 37 to 55% and did not differ significantly between treatment conditions within each experiment (**Supplementary Table 1**).

**Supplementary Table 1.**
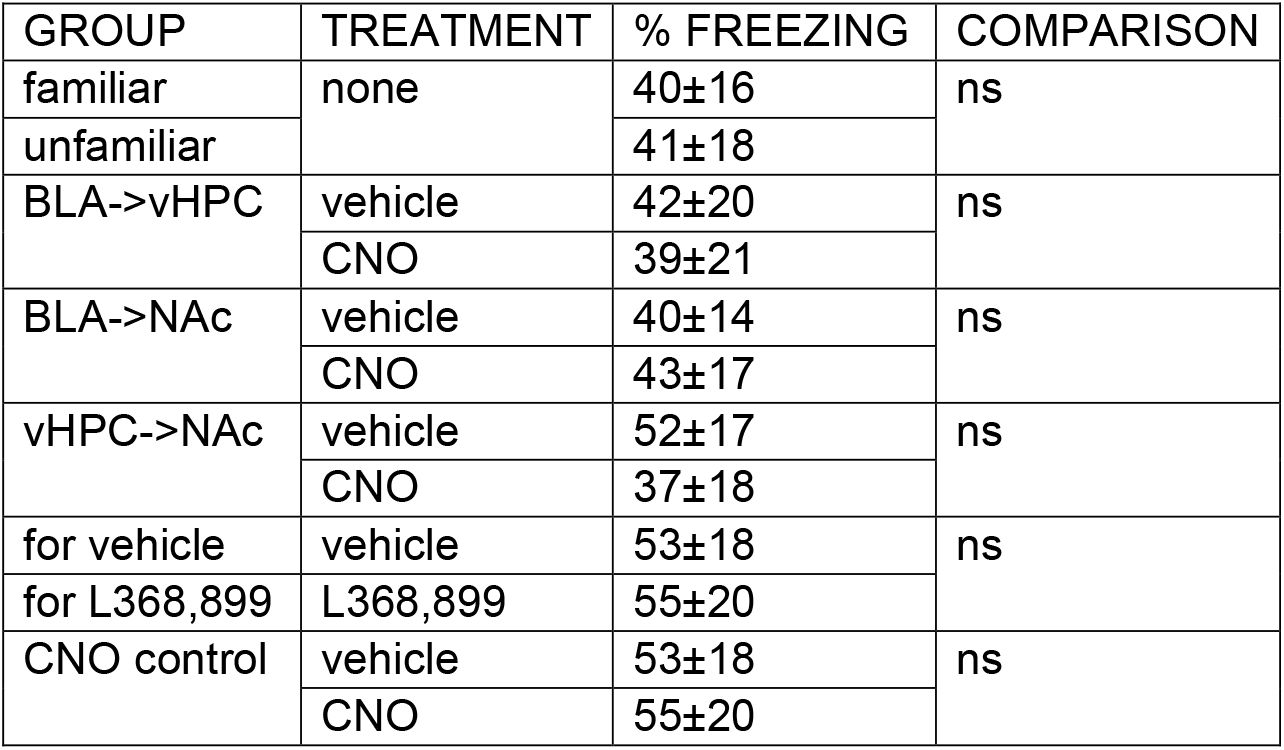
Summary of freezing levels for each experiment.

### Binary viral injections for DREADD inactivation

To selectively express hM4D(Gi) in specific neuronal projections, we used a binary viral strategy. Pseudotyped AAV5-hSyn-DIO-hM4D(Gi)-mCherry (Addgene 44362) was injected at the soma region of interest, and retrograde AAV-hSyn-Cre (Addgene 105553) was injected at the axon terminal region for retrograde transduction. Both AAV stocks were diluted to 5×10^12 viral particles per mL, and 0.2 μL was injected bilaterally at each site. Injection coordinates (AP/ML/DV mm from bregma and brain surface) were as follows: ventral hippocampus (site 1: −3.4/±3.6 /−3.3, and site 2: −3.6/±2.95/−4.5), basolateral amygdala (−1.7/±3.2/−4.3), and nucleus accumbens (+1.4/±0.8/−4.1). Surgical procedures were performed according to established protocols [22].

### Data analysis

Statistical analyses were performed using GraphPad Prism 5 (GraphPad Software, La Jolla, CA). Because CS-induced affiliations were observed in both male and female dyads without sex differences (**Supplementary Fig. 1A**), we combined male and female data for analysis and limited some DREADD experiments to female dyads only (**Fig. 2C, D**). Normality was assessed using the Shapiro–Wilk test. Some datasets exhibited non-normal distributions and were analyzed using the Wilcoxon signed-rank test (paired comparisons) and Mann–Whitney test (unpaired comparisons). All tests were two-sided. Interactions between factors were assessed using two-way repeated-measures ANOVA. Statistical significance was set at p < 0.05.

## Results

### Auditory conditioned stimulus facilitates affiliation in familiar mice

To investigate whether a conditioned stimulus (CS) elicits affiliation, we housed male and female mice as same-sex dyads and subjected them individually to auditory fear conditioning. Dyads were then tested together by exposing them to the CS in a novel context (**Fig. 1A**). Following a 1-minute baseline (pre-CS) period, the CS was presented for 2 minutes.

**Figure 1.**
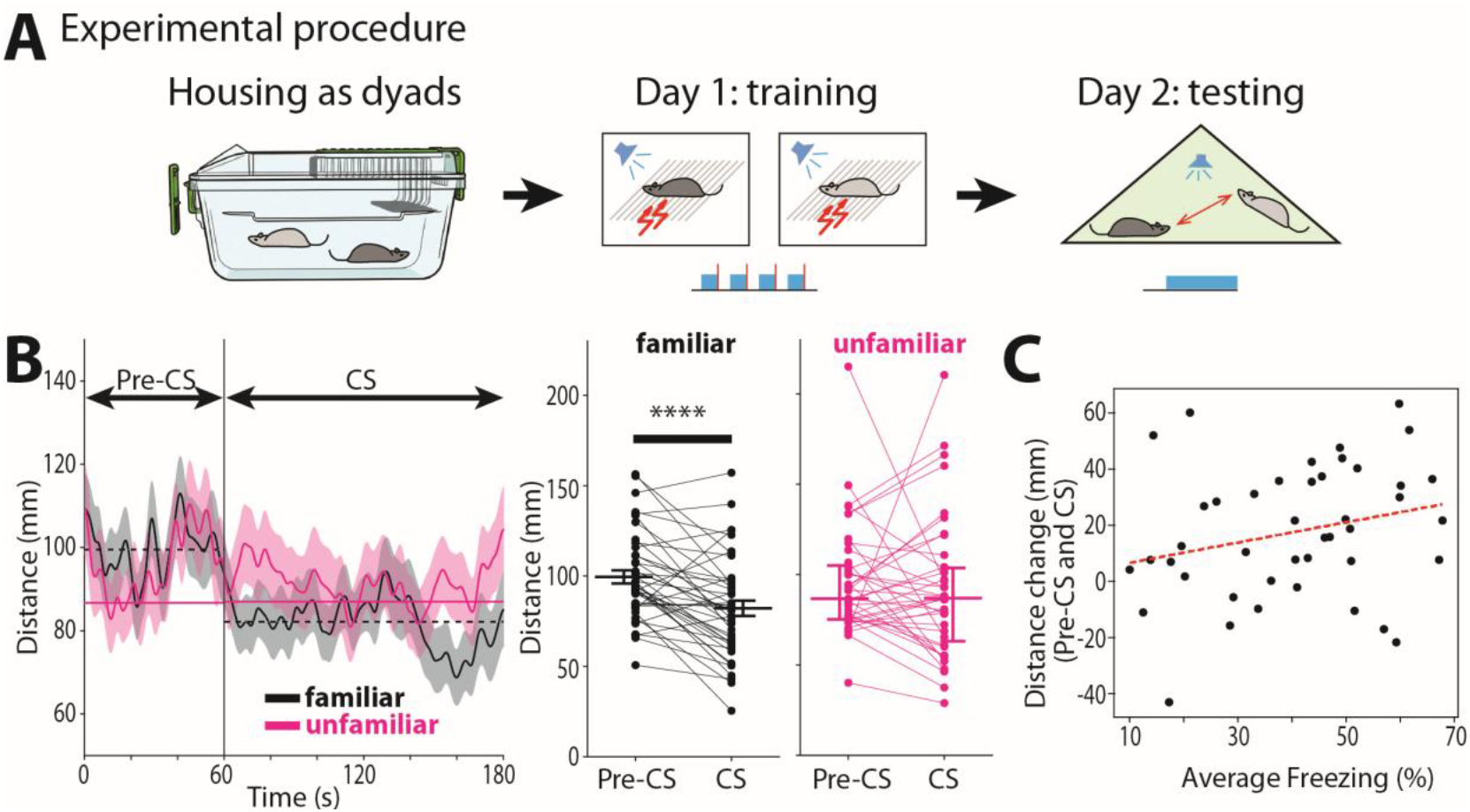
CS-induced social affiliation in dyads requires familiarity and is independent of freezing level. (A) Experimental procedure. (B) Left: dyad distance dynamics throughout pre-CS and CS periods. The solid lines represent mean distances (smoothed by a Gaussian filter, the kernel SD = 5). The shaded areas depict SEMs. The solid and broken horizontal lines represent the medians and means, respectively, for the average dyad distances as analyzed in the paired analyses on the right. Right: average distance during pre-CS and CS periods. **Black:** familiar dyads, n=43 (male:20, female:23), paired t-test: ^****^p<0.0001. **Pink:** unfamiliar dyads, n=39 (male:19, female:20), Wilcoxon Signed Rank test: NS. (C) No correlation between the degree of affiliation (dyadic distance change) and the averaged freezing levels for dyads (Pearson’s coefficient r=0.251, p=0.104).

**Figure 2.**
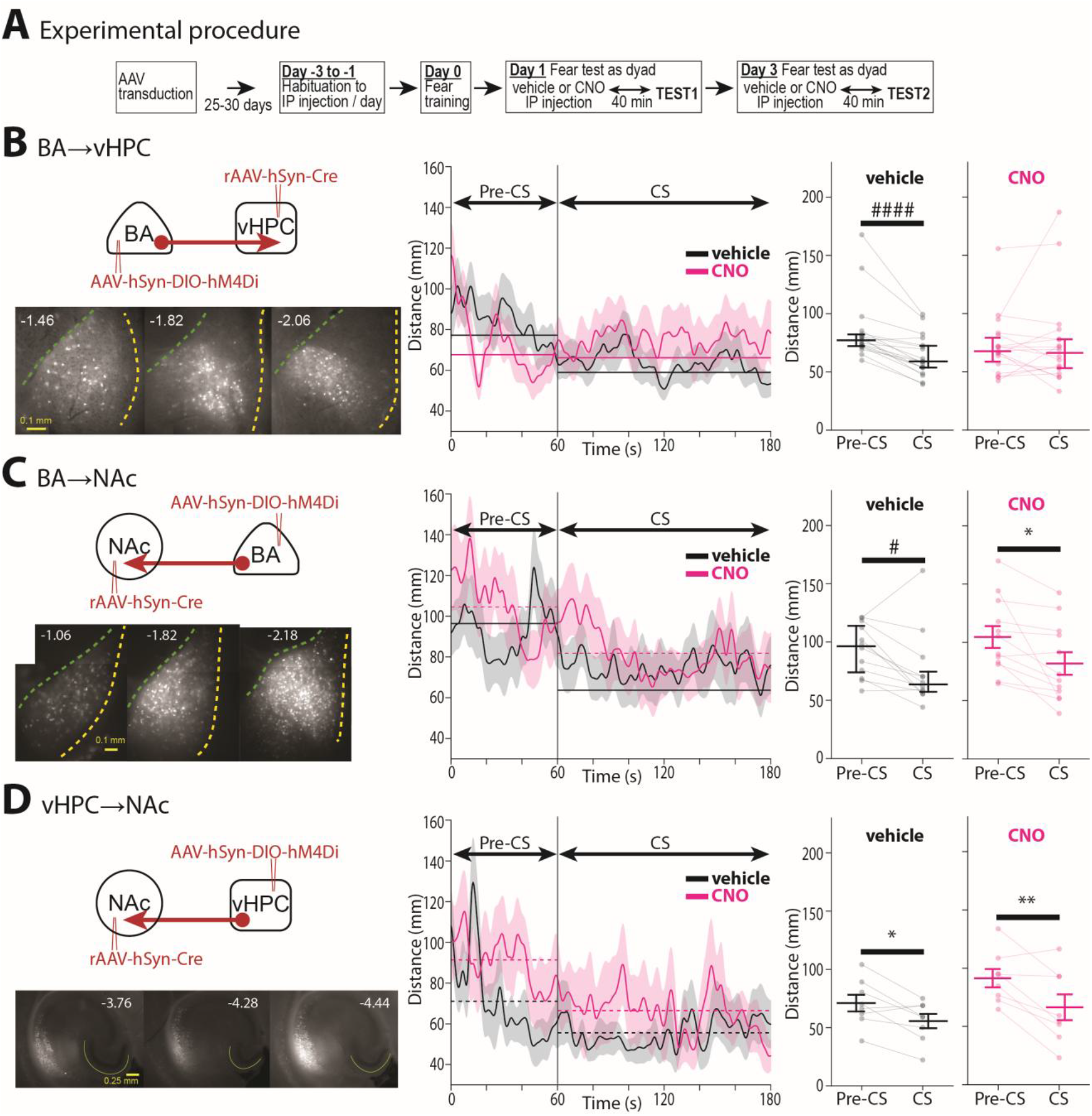
Amygdala-hippocampal projection neurons are required for CS-induced affiliation. (A) Experiment timeline. (B-D) We tested Amygdala-hippocampal (BA➜vHPC) (B), Amygdala-Nucleus Accumbens (BA➜NAc) (C) and Hippocampal-Nucleus Accumbens (vHPC➜NAc) (D) projections in the same way. (Left Upper) Binary strategy to express hM4Di-mCherry in each projection. Red-filled circle and arrow represent hM4Di-mCherry expressing cell body and axon. (Left Lower) mCherry fluorescence in the cell body region, (B,C) BA coronal and (D) vHPC horizontal slices. The yellow and green dashed lines indicate the lateral and medial boundaries of BLA (B and C). The yellow lines mark the dentate gyrus (D). (Center) Distance dynamics throughout the pre-CS and CS periods. (Right) Average distance during pre-CS and CS periods. Details are the same as in **Fig.1B**. (B, BA➜vHPC) n=18 dyads (male:10, female:8), vehicle: ^####^p<0.0001 and CNO: NS (both Wilcoxon signed rank test). (C, BA➜NAc) n=12 female dyads, vehicle: ^#^p<0.05 (Wilcoxon) and CNO: ^*^p<0.05 (Paired t-test). (D, vHPC➜NAc) n=8 female dyads, vehicle: ^*^p<0.05 and CNO: ^**^p<0.01 (both Paired t-test).

We quantified affiliation by measuring snout-to-snout distance. This measure is particularly relevant given that rodent head movements direct gaze [23] and whisker aiming reflects spatial attention [24]. During the CS presentation, snout-to-snout distance decreased significantly compared to the pre-CS period (p<0.0001, **Fig. 1B**), indicating increased proximity between partners. This effect was independent of sex (**Supplementary Fig. 1A**).

To determine whether freezing behavior could account for the observed decrease in distance, we examined the relationship between freezing levels and changes in distance. These variables showed no correlation (**Fig. 1C**), suggesting that independent mechanisms mediate CS-evoked affiliation and freezing. Nevertheless, we summarized freezing levels across all experimental groups in this report in **Supplementary Table 1** and found no significant differences between groups within each experiment.

We next tested whether the distance decrease was a time-dependent process unrelated to CS. A control group of fear-conditioned mice was tested in the absence of CS presentation. These dyads showed no decrease in snout-to-snout distance between the first and last two minutes of the session (**Supplementary Fig. 1B**), demonstrating that CS presentation is necessary for affiliation.

**Supplementary Figure 1.**
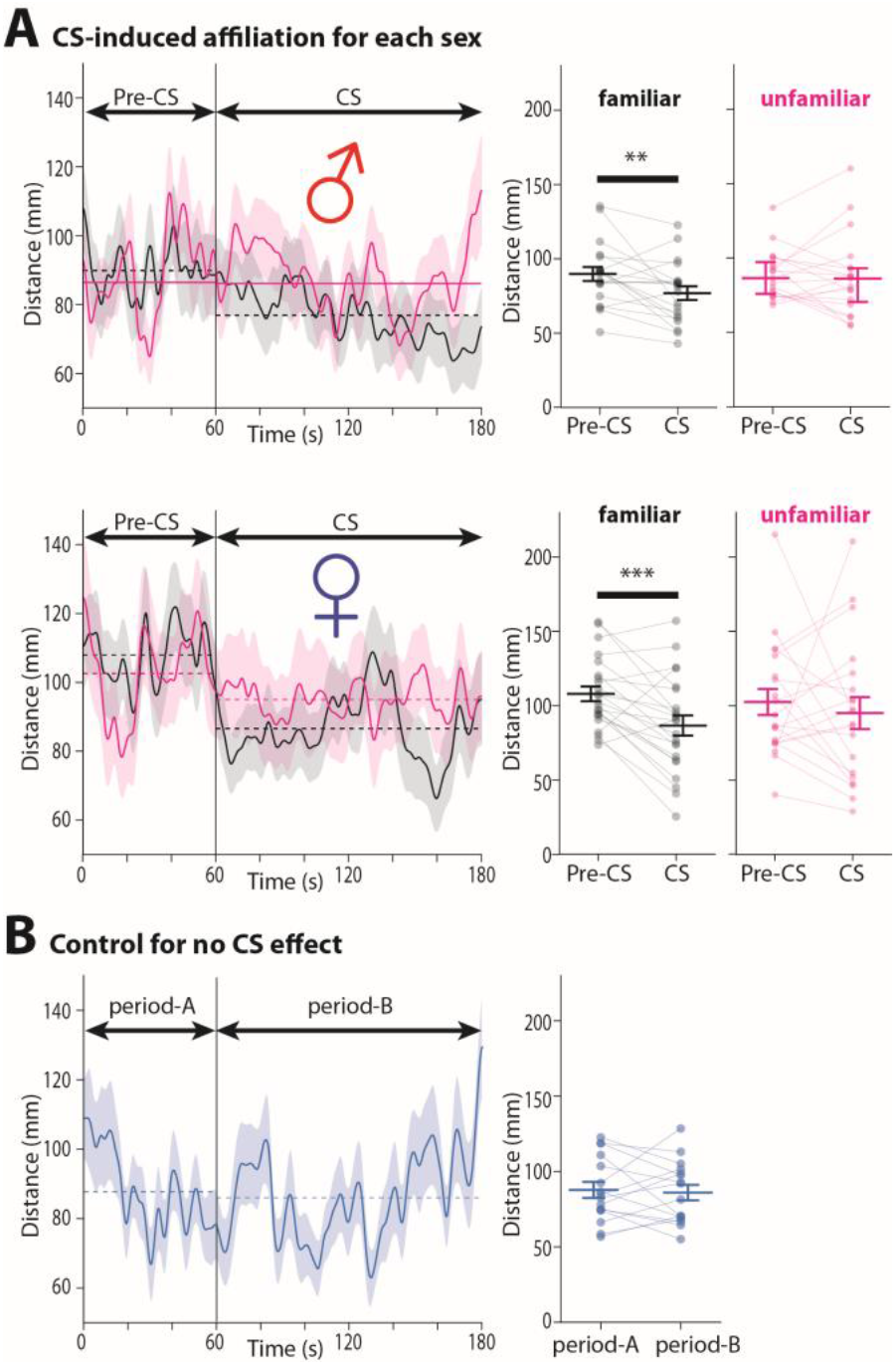
Data split by sex and controls for the experiments in Figure 1. (A) Both sexes exhibit similar CS-induced affiliation. **Upper:** male same-sex dyads (familiar/unfamiliar = 20/19 dyads). **Lower:** female same-sex dyads (familiar/unfamiliar = 23/20 dyads). Details are the same as in Figure 1B. Unfamiliar male dyads: Wilcoxon Signed Rank test: NS. The remaining all groups: Paired t-test: ^**^p<0.01, ^***^p<0.001 and no marking as NS. (B) CS is required for the affiliation. Separate cohort of familiar dyads tested without CS (n=16 dyads, male:9, female:7). Paired t-test: NS.

Finally, we examined whether familiarity between partners influences CS-induced affiliation. When dyads were formed from unfamiliar mice that had not been co-housed, CS presentation failed to decrease snout-to-snout distance (**Fig. 1B, Supplementary Fig. 1A**). This result indicates that familiarity is required for CS-induced affiliation, paralleling findings of a better coordination of innate threat responses among familiar conspecifics in different organisms [25-27].

**Supplementary Figure 2.**
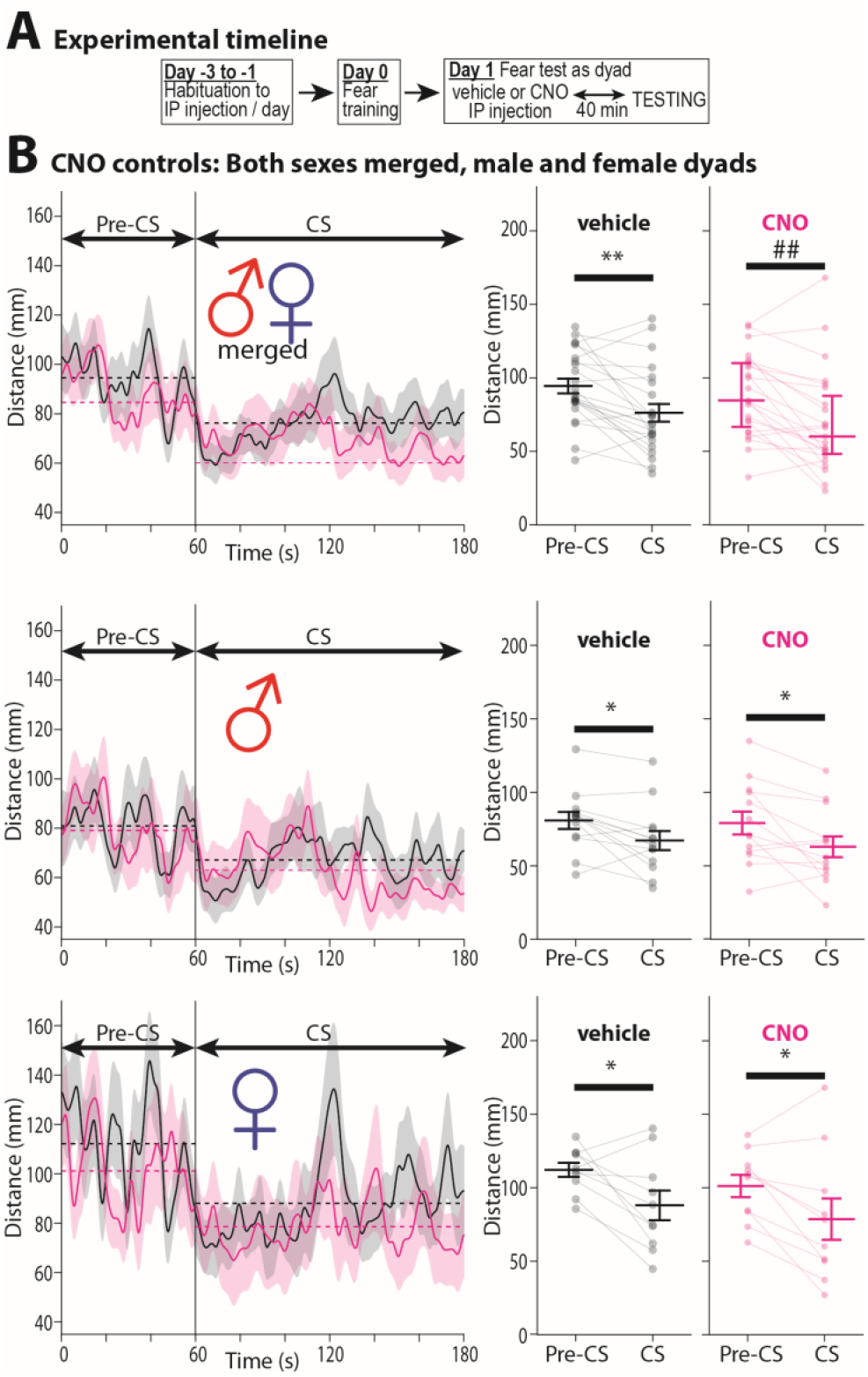
CNO alone does not affect CS-induced affiliation. (A) Experimental timeline. (B) Distance dynamics across the pre-CS and CS periods in merged male and female dyads (n=23). The solid lines represent mean distances smoothed by a Gaussian filter (the kernel SD = 5). The shaded areas depict SEMs. (C) Average distance during pre-CS and CS periods. Data split by sex is shown in D, E (females, n=10) and F, G (males, n=13). Paired t-test: *p<0.05, ^**^p<0.01, Wilcoxon signed rank test: ^##^p<0.01.

### Identification of neuronal circuits required for CS-induced affiliation

The amygdala assigns negative valence to auditory stimuli during fear conditioning and responds to the CS during memory retrieval [28,29]. We therefore hypothesized that CS-activated amygdala neurons drive affiliation through specific downstream targets. Two amygdala projection sites are particularly relevant candidates: the ventral hippocampus (vHPC), which responds to social cues [30] and regulates social approach [31-33], and the nucleus accumbens (NAc), which has been implicated in affiliative behaviors [34-37]. Both structures receive input from the basolateral amygdala (BLA).

To test whether these pathways mediate CS-induced affiliation, we employed chemogenetic inhibition of BLA projections to each target. Before conducting pathway-specific manipulations, we verified that clozapine-N-oxide (CNO) itself does not affect CS-induced affiliation. Control dyads without expressing DREADD effector were injected with either CNO or saline and showed equivalent affiliative responses to the CS (**Supplementary Fig. 2**), ruling out non-specific effects of CNO.

#### BLA➜vHPC Pathway

We first tested the BLA➜vHPC pathway by expressing hM4Di in BLA neurons projecting to vHPC using a binary viral strategy (**Fig. 2B**). Dyads received intraperitoneal injections of either vehicle or CNO before CS testing, with each dyad tested twice under both conditions, separated by a two-day interval (**Fig. 2A**). A two-way repeated-measures ANOVA revealed a significant interaction between the CS and CNO effects on distance (F(1,34)=9.59, p=0.0039). Under vehicle conditions, dyads exhibited robust CS-induced affiliation (p=0.003). In contrast, CNO treatment decreased baseline pre-CS distance (p=0.010, Wilcoxon signed-rank test) and abolished the CS-induced affiliation (**Fig. 2B**). Thus, the BLA➜vHPC pathway contributes to maintaining baseline distance and CS affiliation.

#### BLA➜NAc Pathway

We next tested the BLA➜NAc pathway using the same approach in female dyads (**Fig. 2C**). A two-way repeated-measures ANOVA detected no CS×CNO interaction. Dyads exhibited significant CS-induced affiliation regardless of treatment condition (vehicle: p=0.027; CNO: p=0.011), and CNO did not affect baseline pre-CS distance. These results indicate that the BLA-to-NAc pathway is not required for CS-induced affiliation.

#### vHPC➜NAc Pathway

Given that the vHPC➜NAc pathway regulates approach/avoidance behaviors toward conspecifics [31,32] and we confirmed that the BLA➜vHPC Pathway influences intermouse distance (**Fig. 2B**), we tested the role of vHPC neurons projecting to NAc using the same chemogenetic approach in female dyads (**Fig. 2D**). A two-way repeated-measures ANOVA detected no CS×CNO interaction. Dyads exhibited significant CS-induced affiliation under both vehicle (p=0.039) and CNO (p=0.023) conditions. However, CNO treatment increased baseline pre-CS distance relative to vehicle (p=0.022, Paired t-test; **Fig. 2D**). Thus, the vHPC➜NAc Pathway contributes to baseline distance but does not to CS-induced affiliation.

### Oxytocin signaling is required for CS-induced affiliation

Oxytocin signaling increases the salience of social stimuli [38] and supports social behaviors [39]. We therefore tested whether oxytocin signaling is required for CS-induced affiliation by administering the oxytocin receptor antagonist L368,899 (3 mg/kg, intraperitoneal) or saline 1 hour before CS testing. This dose was selected based on previous studies demonstrating modulation of social behaviors [40-42]. Each dyad was tested only once under either antagonist or vehicle conditions (**Fig. 3A**).

**Figure 3.**
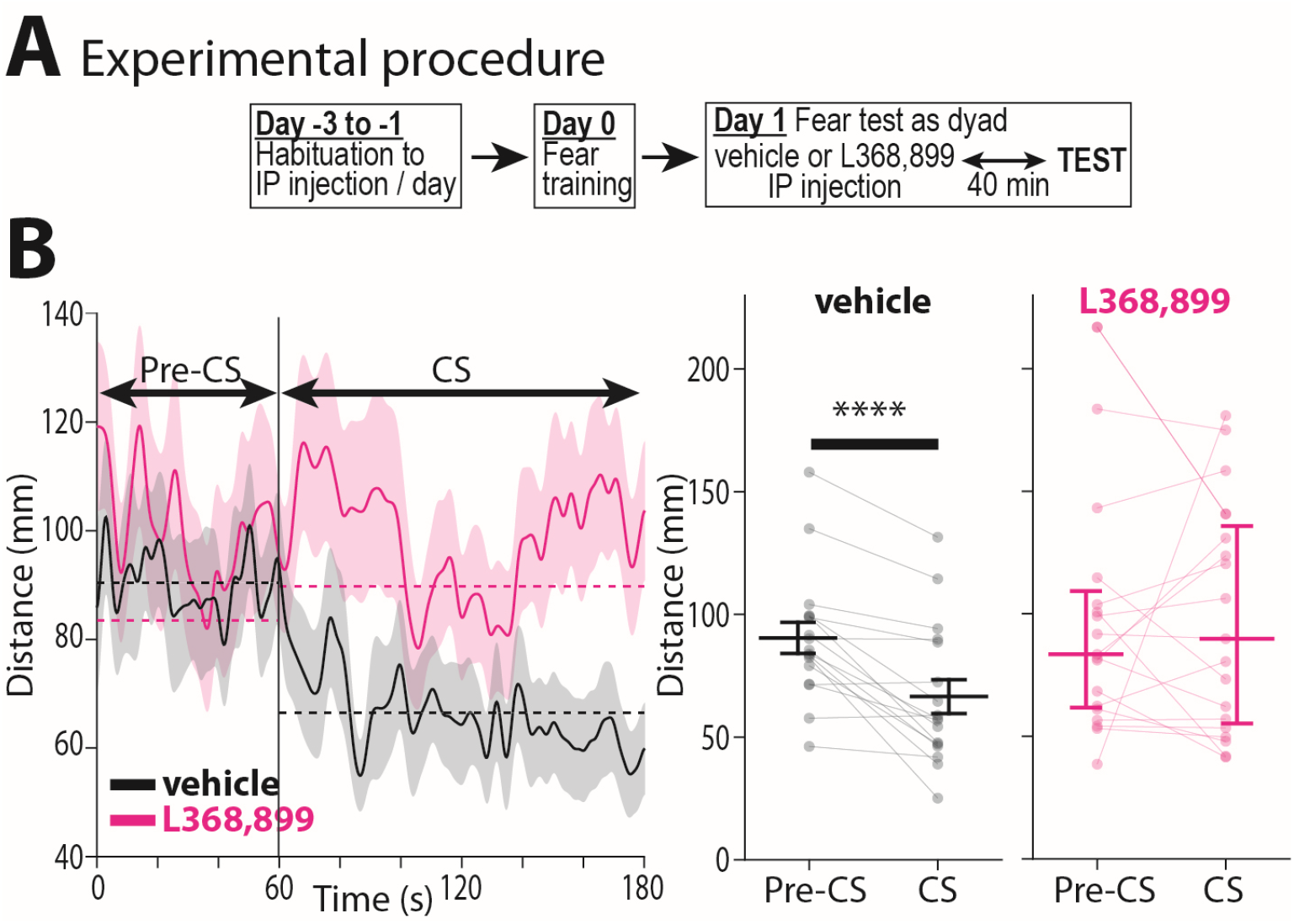
Oxytocin receptor antagonist L368,899 prevents CS-induced affiliation. (A) Experimental timeline. (B) Distance dynamics throughout the pre-CS and CS periods. (Right) Average distance during pre-CS and CS periods. Details are the same as in **Fig.1B**. Vehicle: n=17 dyads (male:9, female:8), ^****^p<0.0001 (Paired t-test). L368,899: n=19 dyads (male:9, female:10), no marking as NS (Wilcoxon signed rank test).

L368,899 did not affect baseline intermouse distance during the pre-CS period (p>0.05, Mann-Whitney U test). A mixed-model ANOVA revealed a CS×treatment interaction approaching significance (F(1,34)=3.81, p=0.06). In the saline-treated group, CS presentation significantly decreased intermouse distance (p=0.0001), whereas the antagonist-treated group showed no change in intermouse distance during CS (p>0.05). These results demonstrate that oxytocin receptor signaling is necessary for CS-induced affiliation.

## Discussion

This study demonstrates that the auditory conditioned fear response includes not only freezing but also affiliation with a familiar conspecific. This CS-induced affiliation occurs independently of freezing and requires the BLA➜vHPC pathway and oxytocin receptor signaling.

Defensive aggregation is a widespread phenomenon in which threatened animals reduce individual risk by decreasing interindividual distance [43]. While defensive aggregation has been extensively documented in response to innate threat stimuli, particularly predator-related cues [7-15], our findings demonstrate for the first time that a learned threat signal—an auditory CS—can also elicit proximity-seeking in same-sex dyads of laboratory mice. This finding reveals that the conditioned fear response encompasses both non-social defensive behaviors, such as freezing, and social approach behaviors. However, these two types of defensive responses occur independently of each other, as evidenced by the absence of correlation between them and the lack of freezing alterations across all circuit manipulations (**Supplementary Table 1**). This independence suggests that CS-activated freezing pathways, which include connections from the lateral to the central amygdala [44,45], operate independently of CS-activated affiliation circuits.

The requirement for familiarity in CS-induced affiliation implicates social recognition systems, including the hippocampus, which mediates social memory [32,46]. Chemogenetic inhibition of the BLA➜vHPC pathway abolished CS-induced affiliation without altering freezing, consistent with a model in which the amygdala detects the threatening CS and transmits this information to the hippocampus to drive affiliation. However, this manipulation also increased baseline proximity during the pre-CS period, suggesting that tonic BLA➜vHPC activity suppresses affiliation in the absence of threat. This interpretation aligns with findings that optogenetic inhibition of BLA➜vHPC projections increases social approach [47]. These observations argue against a simple model in which “CS-activated amygdala drives affiliation.” Instead, CS presentation may suppress a subset of BLA neurons projecting to vHPC, thereby disinhibiting affiliative behavior. This interpretation is supported by evidence that CS upregulates some BLA neurons while downregulating others [28], and that BLA sends both inhibitory and excitatory projections to vHPC with potentially distinct CS responses [48]. Future studies monitoring BLA➜vHPC projection dynamics during CS presentation will be necessary to test these possibilities.

Inhibition of BLA➜NAc projections did not affect either CS-induced affiliation or baseline intermouse distance, which was unexpected given evidence that the NAc regulates social approach [37,49,50] and BLA inputs to NAc regulate sociability [51]. In contrast, inhibiting vHPC➜NAc projections increased baseline intermouse distance, consistent with reports that vHPC➜NAc pathway activity correlates with social interaction in mice [52] and that its inhibition decreases social interaction in rats [31]. However, this manipulation did not prevent CS-induced affiliation, suggesting that CS recruits a parallel pathway that bypasses or overrides this circuit. The vHPC projections to the lateral septum are promising candidates, given their roles in regulating approach-avoidance decisions toward novel conspecifics [33] and in motivational conflict [53], and the finding that rats showing greater predator odor-induced aggregation exhibit elevated c-Fos expression in the lateral septum [17].

We tested the role of oxytocin signaling using systemic administration of the high-affinity oxytocin receptor antagonist L-368,899, which crosses the blood-brain barrier [41] and modulates social behaviors in mice [40-42]. The antagonist abolished CS-induced affiliation, consistent with evidence across species that oxytocin and its homologs enhance affiliative behaviors, including flocking in birds [54], shoaling in fish [55] and predator-odor-induced aggregation in rats [18]. Studies in prairie voles and California mice have identified the NAc, anterior cingulate cortex, and BLA as sites where oxytocin regulates social approach [37,56]. Whether the oxytocin signaling required for CS-induced affiliation acts within the BLA-to-vHPC circuit or at other sites remains to be determined through circuit-specific manipulations of oxytocin receptor signaling.

## Data and code availability

All primary data, including video files, are available from the authors upon reasonable request. Analysis code and example datasets will be provided as part of a replication package upon publication.

## Acknowledgements

This work was supported by the National Institutes of Health Grant No R21MH137592 and the Seale Innovation Fund to AM.The authors thank Ayush Pinnamaraju for manual annotation of mouse behaviors in videos and for verifying the expression of DREADD effectors in the tested mice. The authors report no biomedical financial interests or potential conflicts of interest.

## Author Contributions

WI and AM conceived and designed the study. AM performed the experiments. WI wrote custom code for data analysis. WI and AM analyzed the data. AM wrote the initial draft of the manuscript, which was refined by WI and AM.

