## Supplemental Figures 1,2 and Table for "Affiliation in mice induced by an auditory-conditioned stimulus"

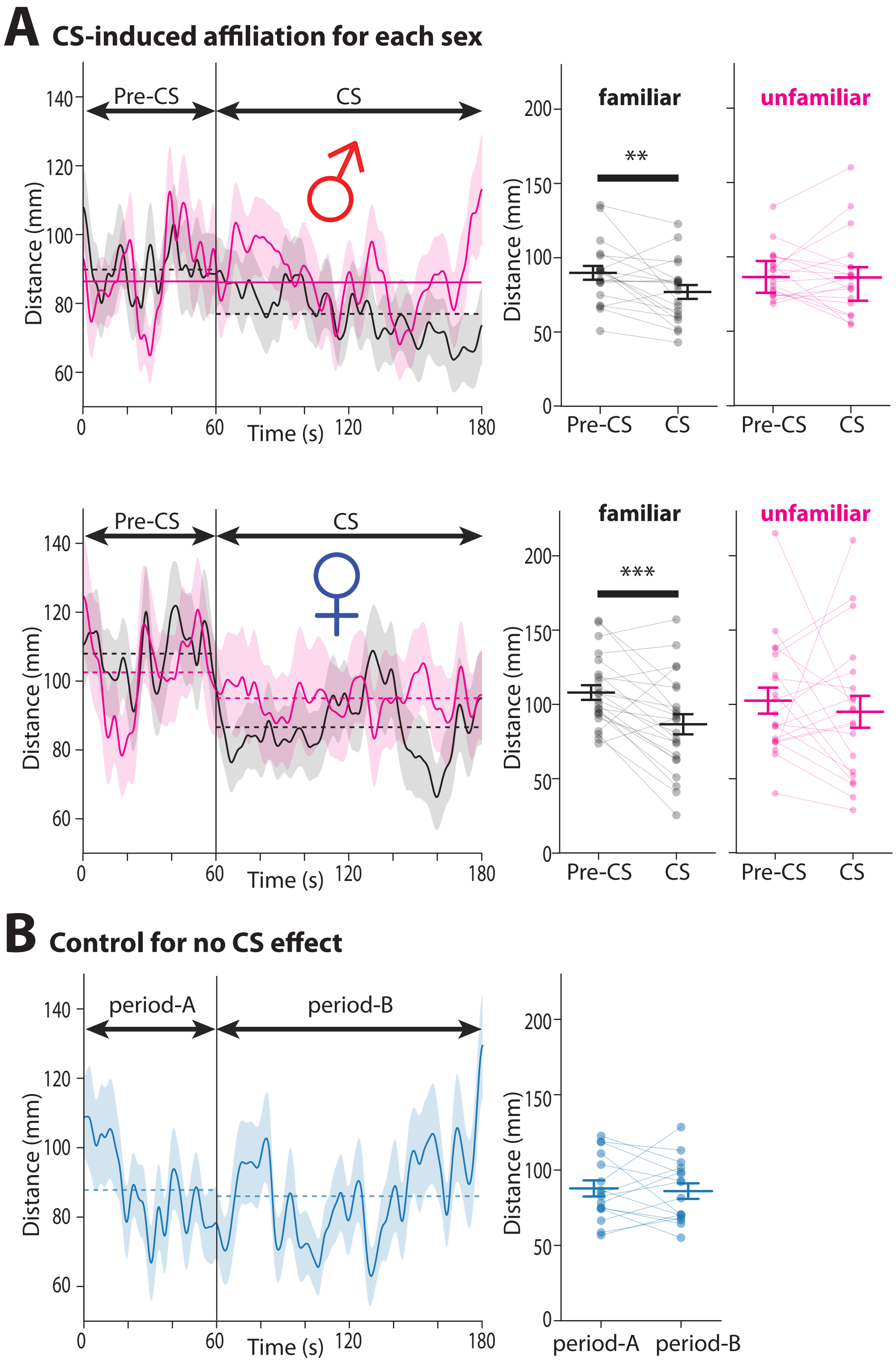


**Supplementary Figure 1. Data split by sex and controls for the experiments in Figure 1.** (A) Both sexes exhibit similar CS-induced affiliation. **Upper:** male same-sex dyads (familiar/unfamiliar = 20/19 dyads). **Lower:** female same-sex dyads (familiar/unfamiliar = 23/20 dyads). Details are the same as in Figure 1B. Unfamiliar male dyads: Wilcoxon Signed Rank test: NS. The remaining all groups: Paired t-test: $^{**}p<0.01$, $^{***}p<0.001$ and no marking as NS. (B) CS is required for the affiliation. Separate cohort of familiar dyads tested without CS (n=16 dyads, male:9, female:7). Paired t-test: NS.

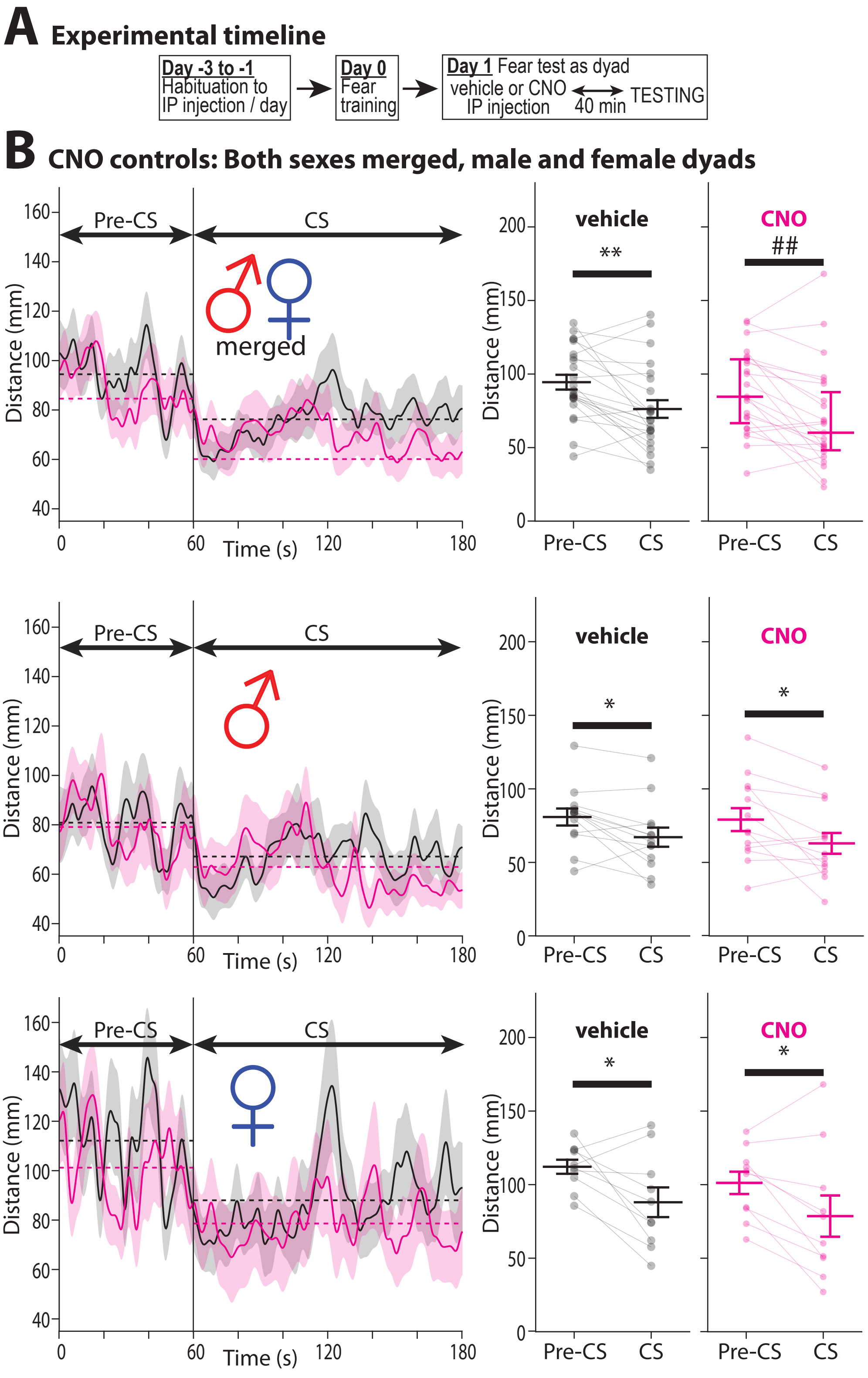


**Supplementary Figure 2**. **CNO alone does not affect CS-induced affiliation.** (A) Experimental timeline. (B) Distance dynamics across the pre-CS and CS periods in merged male and female dyads (n=23). The solid lines represent mean distances smoothed by a Gaussian filter (the kernel SD = 5). The shaded areas depict SEMs. (C) Average distance during pre-CS and CS periods. Data split by sex is shown in D, E (females, n=10) and F, G (males, n=13). Paired t-test: *$p<0.05$, **$p<0.01$, Wilcoxon signed rank test: ##$p<0.01$.

**Supplementary Table 1.** Summary of freezing levels for each experiment.

| GROUP | TREATMENT | % FREEZING | COMPARISON |
|---|---|---|---|
| familiar | none | 40±16 | ns |
| unfamiliar | | 41±18 | |
| BLA->vHPC | vehicle | 42±20 | ns |
| | CNO | 39±21 | |
| BLA->NAc | vehicle | 40±14 | ns |
| | CNO | 43±17 | |
| vHPC->NAc | vehicle | 52±17 | ns |
| | CNO | 37±18 | |
| for vehicle | vehicle | 53±18 | ns |
| for L368,899 | L368,899 | 55±20 | |
| CNO control | vehicle | 53±18 | ns |
| | CNO | 55±20 | |
